# Multiplexed nanopore sequencing of HLA-B locus in Māori and Polynesian samples

**DOI:** 10.1101/169078

**Authors:** K.N.T. Ton, S.L. Cree, S.J. Gronert-Sum, T.R. Merriman, L.K. Stamp, M.A Kennedy

## Abstract

The human leukocyte antigen (HLA) system is a gene family that encodes the human major histocompatibility complex (MHC). HLA-B is the most polymorphic gene in the MHC class I region, comprised of 4,765 HLA-B alleles (IPD-IMGT/HLA Database Release 3.28). Many HLA-B alleles have been associated with adverse drug reactions and disease risks, and we are interested in developing efficient methods for analysis of HLA alleles in this context. Here we describe an approach to HLA-B typing using multiplexed next generation sequencing on the MinION™ nanopore sequencer (Oxford Nanopore Technologies), combined with data analysis with the SeqNext-HLA software package (JSI Medical Systems GmbH, Ettenheim, Germany). The nanopore sequencer offers the advantages of long-read capability and single molecule reads, which can facilitate effective haplotyping. We developed this method using reference samples of known HLA-B type as well as individuals of New Zealand Māori or Pacific Island (Polynesian) descent, because HLA-B diversity in these populations is not well understood. We demonstrate here that nanopore sequencing of barcoded, pooled, 943 bp polymerase chain reaction (PCR) amplicons of 49 DNA samples, on one R9.4 flowcell (Oxford Nanopore Technologies), generated ample read depth for all samples. Sequence analysis using SeqNext-HLA software assigned HLA-B alleles to all samples at high-resolution with very little ambiguity. Our PCR-based next generation sequencing method is a scaleable and efficient approach for genotyping HLA-B and potentially any other HLA locus. Finally, we report our findings on HLA-B genotypes of this cohort, which adds to our understanding of HLA-B allele frequencies among Māori and Polynesian people.

## Introduction

The human leukocyte antigen (HLA) locus contains a large family of genes encoding the human major histocompatibility complex (MHC) proteins. It is located on chromosome 6p21 and divided into three classes: class I, class II, and class III. The primary function of HLA genes is to present peptide antigens to T-cells in the adaptive immune system. HLA molecules are extremely variable due to their peptide-binding function and are associated with autoimmune diseases and adverse drug reactions (Tiwari & Terasaki, 2012, Bharadwaj *et al.*, 2010). HLA-B is the most polymorphic gene, with over 4,600 known alleles encoding 3,408 unique proteins (IMGT/HLA Database release 3.27 in January 2017) (Robinson *et al.*, 2014). Previous studies have identified particular HLA-B alleles as risk factors for drug-induced hypersensitivity reactions (Alfirevic & Pirmohamed, 2010, Sukasem *et al.*, 2014). For example, screening for the HLA-B∗57:01 allele is recommended prior to abacavir treatment to decrease risk of a hypersensitivity reaction. (Martin *et al.*, 2012). Strong association between HLA-B∗58:01 and allopurinol-induced severe cutaneous adverse reactions such as Stevens–Johnson syndrome (SJS) or toxic epidermal necrolysis (TEN) have been reported across different populations (Somkrua *et al.*, 2011). Similarly, HLA-B∗15:02 has been associated with carbamazepine and phenytoin-induced SJS or TEN in Asian patients (Yip *et al.*, 2012).

The high level of polymorphism in the MHC family means HLA genotyping is complex. HLA alleles are mostly determined by the sequences of exons 2 and 3 in HLA class I genes and exon 2 in HLA class II genes (Shiina *et al.*, 2012). Present DNA-based methods for HLA typing are polymerase chain reaction (PCR) -sequence-specific priming (PCR-SSP), PCR-sequence-specific oligo hybridization (PCR-SSO), PCR-restriction fragment length polymorphism (PCR-RFLP), and sequence-based typing (SBT) (Erlich, 2012, Bontadini, 2012, Tait *et al.*, 2009). SBT is currently considered the gold standard method applied in high-resolution HLA typing (Erlich, 2012). However, this approach may generate ambiguous HLA typing due to haplotype phase issues and incomplete sequencing. Other approaches have various limitations in resolution, workflow complexity, and probe design and testing requirements, as new HLA alleles are submitted to the IMGT/HLA sequence database (Erlich, 2012, Shiina *et al.*, 2012). Recently, next-generation sequencing (NGS) methods have become widely established for HLA typing (Bentley *et al.*, 2009, Abbott *et al.*, 2006, Erlich, 2012, Erlich *et al.*, 2011, Hosomichi *et al.*, 2013, Shiina *et al.*, 2012, Hosomichi *et al.*, 2015, Schöfl *et al.*, 2017). Such approaches reduce the risk of phase ambiguity and allow high-throughput, high-resolution HLA typing. Various approaches to HLA typing using NGS have been developed, including PCR-based HLA sequencing (Boegel *et al.*, 2012, Erlich *et al.*, 2011, Hosomichi *et al.*, 2013, Liu *et al.*, 2012, Shiina *et al.*, 2012, Schöfl *et al.*, 2017), whole exome sequencing (WES) or whole genome sequencing (WGS) data-derived typing (Liu *et al.*, 2012, Major *et al.*, 2013).

Here we describe (a) the development of high-throughput HLA typing from next-generation DNA sequencing data, focusing on the HLA-B locus, and (b) its application to identifying HLA-B alleles within the Māori and Polynesian population of New Zealand. Our strategy took advantage of a recent iteration of the novel NGS platform, the MinION™ nanopore sequencer (Oxford Nanopore Technologies), and barcode sequences for labeling and simultaneously analysing HLA-B amplicons from 49 samples. The MinION is a tiny, portable nanopore sequencer powered by a USB 3.0 port (Quick *et al.*, 2014). It allows analysis of sequencing data in real time and generation of very long reads (Urban *et al.*, 2015). A small pilot study used the device to examine HLA-A and HLA-B alleles from a single sample, using the earlier R7.0 flowcell chemistry, but that study failed to call HLA haplotypes due to the high error rates of the device at that time (Ammar *et al.*, 2015). Oxford Technologies released a new chemistry (R9.4) in May 2016, which was expected to be faster, more accurate and with higher throughput. This major update motivated us to examine the performance of this pocket-sized device on one of the most polymorphic genes in human body, HLA-B.

To date, there is a paucity of studies providing prevalence data of HLA-B alleles in Māori and Polynesian population in New Zealand (Abbott *et al.*, 2006, Edinur *et al.*, 2013). Given that HLA-B alleles are so relevant to disease predisposition and adverse drug reactions, it is important to establish a prevalence dataset for HLA-B for these population groups.

## Methods

### Participants

Forty unrelated individuals with no history of inflammatory disorders were recruited from the Otago and Auckland regions of New Zealand. All participants gave their written informed consent. Genomic DNA was prepared by guanidinium-HCl based chloroform extraction. Additionally, five individuals were included from a local study on adverse drug reactions called <u>U</u>nderstanding adverse <u>D</u>rug Reactions or responses <u>U</u>sing <u>G</u>enome <u>S</u>equencing (UDRUGS), and samples from four individuals of known HLA-B genotype were obtained from the Coriell Institute for Medical Research (Camden, NJ, USA). These nine individuals were used as a reference set for the MinION analysis, after confirmation by either Sanger sequence based typing (below) or data retrieved from the 1000 Genomes project, or both.

### HLA-B genotyping by Sanger sequencing

We selected a subset of our participants (four Polynesian, five UDRUGS and two Coriell samples) to analyze by Sanger sequencing, as additional references. Nested PCR was used to amplify a 1710 bp region spanning exon 2 and exon 3 of HLA-B. PCR products were diluted and used as templates in second round PCR to amplify a 943 bp amplicon. These amplicons were then directly sequenced in both forward and reverse directions using a set of six sequencing primers (Table 1). The primers used for amplification included some nucleotide redundancies at sites of HLA-B variation, to prevent allelic drop-out during the PCR step. All PCR primers and sequencing primers were derived from published work (Abbott *et al.*, 2006, Cotton *et al.*, 2012). The HLA-B genotypes of these 11 samples were generated from the Sanger sequence data using SBTengine v.3.10.0.2610 (GenDX, Utrecht, Holland).

**Table 1.**
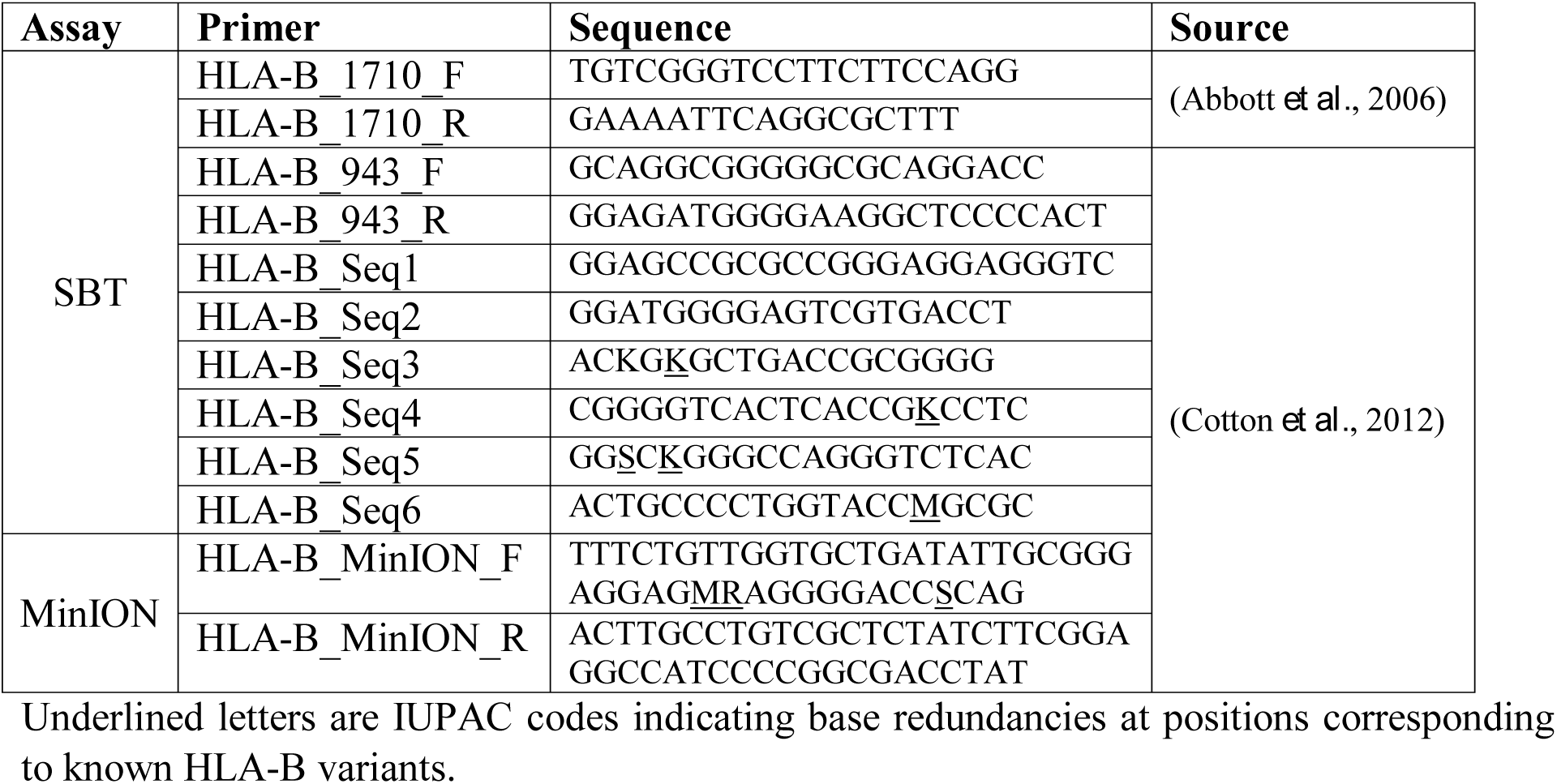
Primers used for amplifying and sequencing

### MinION library construction

The primer used to amplify a fragment of 943 bp HLA-B exon 2 and 3 included a specific sequence (Table 1) at the 5’ end, which is compatible with barcode sequences (Oxford Nanopore Technologies). A standard protocol of the Kapa LongRange Hotstart DNA PCR (Kapa Biosystems) was applied, consisting of 1X Kapa LongRange Buffer (without Mg^2+^), 2.0 mM MgCl_2_, 0.3 mM dNTPs (10mM each dNTP), 0.5 μM of each primer, 1.25 U/50 μl Kapa LongRange HotStart DNA Polymerase, 50 ng genomic DNA, and water up to 50 μl. Thermal cycling conditions were 94°C for 3 min, 25 cycles at 94°C for 15 s, 68°C for 15 s, and 72°C for 1 min, with a final extension at 72°C for 1 min. The PCR products were visualized by electrophoresis on 2% agarose gel stained with SYBR™ Safe DNA Gel Stain (Invitrogen), and then purified using 1x Agencourt AMPure XP beads (Beckman Coulter).

PCR products were quantified by Qubit^®^ 2.0 Fluorometer (ThermoFisher Scientific) and were diluted to 2 nM in water. A second PCR was performed to incorporate barcode sequences using Oxford Nanopore PCR Barcoding kit (EXP-PBC096). Each 100 μl reaction contained 1X Kapa LongRange Buffer (without Mg^2+^), 2.0 mM MgCl_2_, 0.3 mM dNTPs (10mM each dNTP), 0.2 μM PCR Barcode primers (from BC01 to BC49), 2.0 U Kapa LongRange HotStart DNA Polymerase, and 0.5 nM of first-round PCR product. The cycling parameters were an initial denaturation 95°C for 3 min, followed by15 cycles at 95°C for 15 s, 62°C for 15 s, and 65°C for 1 min, with a final extension at 65°C for 1 min. All 49 barcoded products were cleaned up with 1x Agencourt AMPure XP beads, then quantified. Purified PCR products were normalized by concentration before being pooled for library preparation.

The pooled library was prepared using the Oxford Nanopore Sequencing protocol (SQK-NSK007). We used 5 μg of library as an input, instead of the recommended 1 μg, to improve yield for downstream steps. Briefly, 5 μg of purified amplicon library was prepared with the NEBNext end repair module (New England Biolabs), then dA-tailed using the NEBNext dA-tailing module (New England Biolabs). The end-prepared, dA-tailed library was subsequently ligated with leader and hairpin adapters, followed by purification using Dynabeads^®^ MyOne™ Streptavidin C1 beads (Invitrogen).

The final prepared library from 49 participants was loaded into the MinION R9.4 flowcell (Oxford Nanopore Technologies). The flowcell was run for 48 hours using the MinKNOW software (0.51.1.39).

### Data analysis

Raw sequence data were uploaded for base-calling using Metrichor software (2D Basecalling for SQKMAP007 - v1.107). Sequences in FASTA format were extracted from the raw FAST5 files using poretools v.0.6.0 (Loman & Quinlan, 2014). We only used two-dimensional (2D) reads, which are consensus calls of the combined template and complement strands, to perform HLA-B locus high-resolution typing with SeqPilot v.4.3.1 (JSI medical systems). HLA-B ambiguities were designated as G group nomenclature (http://hla.alleles.org/alleles/g_groups.html). If SeqPilot software could not call the genotype without mismatch, we checked all best matched alleles and manually assigned them.

## Results

A total of 40 Polynesian, four Coriell and five UDRUGS individuals were selected for library construction. We successfully amplified a region of 943 bp spanning exon 2 and 3 of HLA-B, using long-range PCR. After purifying, all 49 PCR products were diluted to reach the desired concentration (2.0 nM). Three gave insufficient yield, ranging from 0.19 to 1.47 nM. However, these three were still included to test whether they could still be effectively amplified in the barcoding step.

Two primers used in the first PCR were tailed with the adapter sequences, which were compatible with Oxford Nanopore barcodes. Each PCR product was then amplified with barcode primers, at which point all 49 PCR products were tagged with barcodes, increasing their length to 1063 bp (Figure 1).

**Figure 1.**
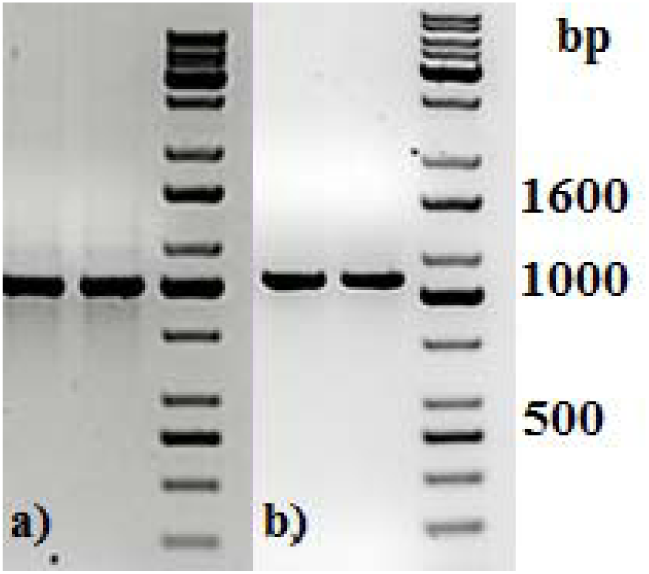
Agarose gel electrophoresis of two representative long-range PCR amplicons (a) before and (b) after PCR barcoding step. DNA molecular size marker obtained from KAPA Universal Ladder (KAPA Biosystems, Boston, MA).

PCR products were subjected to normalization prior to pooling and sequencing on the MinION (~368 ng/each). Five samples had significantly lower concentrations than other samples (range: 20.4 – 66.8 ng), but these were included in the pool (Table 2). The total DNA quantity in the pool was 7.5 μg and 5 μg was used for downstream steps. After end-repair, adapter ligation and purification steps, 585 ng of prepared library remained and was loaded into the MinION flow cell.

**Table 2.** DNA quantity and number of mapped reads of poorly amplified samples

| Individual | Index | DNA quantity (ng) | Mapped 2D read (number) |
| --- | --- | --- | --- |
| PI_G2 | BC15 | 66.84 | 473 |
| PI_C3 | BC19 | 20.4 | 80 |
| PI_D3 | BC20 | 57.96 | 869 |
| PI_H3 | BC22 | 64.44 | 333 |
| PI_C6 | BC42 | 33.36 | 158 |

Given that for this version of the MinION chemistry, 2D reads were more accurate and had greater length than 1D reads, we extracted only the 2D reads for downstream analysis. After conversion, all 49 FASTQ files were imported into SeqPilot software for HLA-B allele assignment. The mean read length was 1029 bp, close to the expected size of all amplicons (Figure 2).

**Figure 2.**
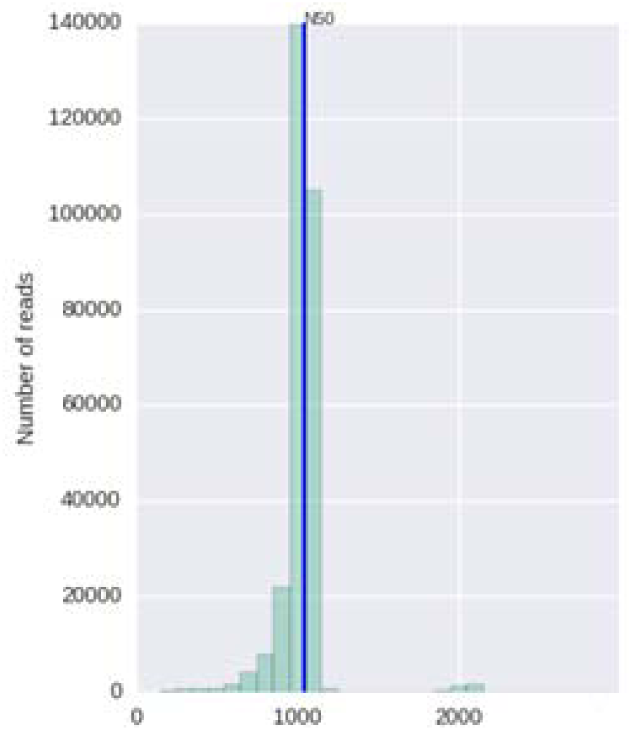
Histogram of read length with read N50 metric

An average of 5,854 sequence reads per barcoded sample was obtained from a total of 289,095 2D reads. There were 286,852 reads parsed to uniquely identified barcodes, of which 199,297 reads correctly aligned to the assigned allele sequences. The distribution histogram of both assigned and aligned reads for each barcoded sample is shown in Figure 3.

**Figure 3.**
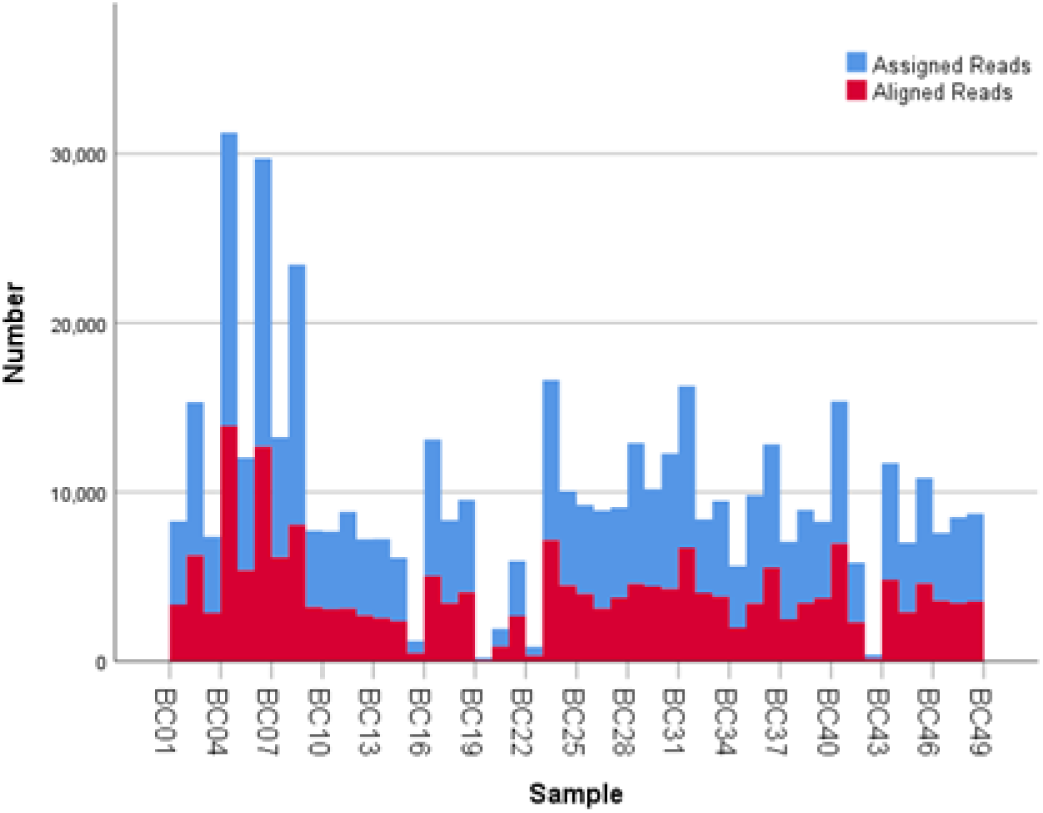
Number of mapped and unmapped reads per individual

**Figure 3.**
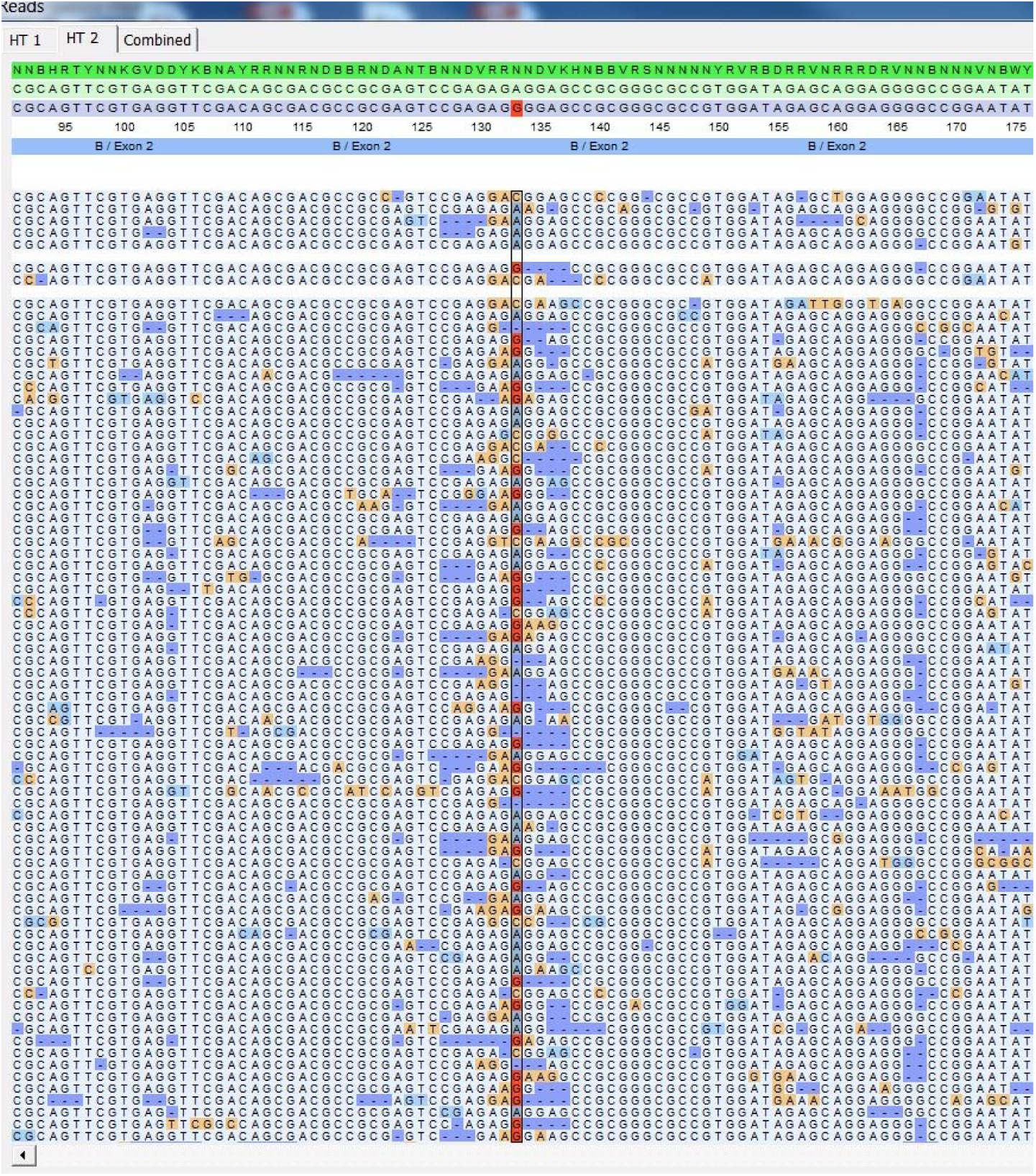
SeqPilot screenshot from mapping sequencing reads with reference sequences. Bias phasing position (red) followed by deletion errors.

The five samples that amplified poorly still produced ample reads for SeqPilot to generate HLA-B typing calls (Table 2). Notably, individual PI_C3 only had 80 reads that aligned to the reference sequence and was HLA-B∗44:04/56:02:01. We are confident these alleles are correct as no mismatches occurred at key polymorphic sites in the reads.

Using FASTQ files as inputs, SeqPilot effectively assigned HLA-B alleles of 49 individuals. There were 4 homozygotes and 45 heterozygotes, resulting in 38 alleles called at 4-digit resolution and five alleles at 2-digit resolution (Table 3). There were six that could not be automatically called by the software due to mismatches with reference sequences. In this case, we had to visually inspect the mismatched sites in the SeqPilot interface, and manually re-phased bases. For these six, we manually assigned allele pairs from the suggested list, choosing those with the least mismatch.

**Table 3.**
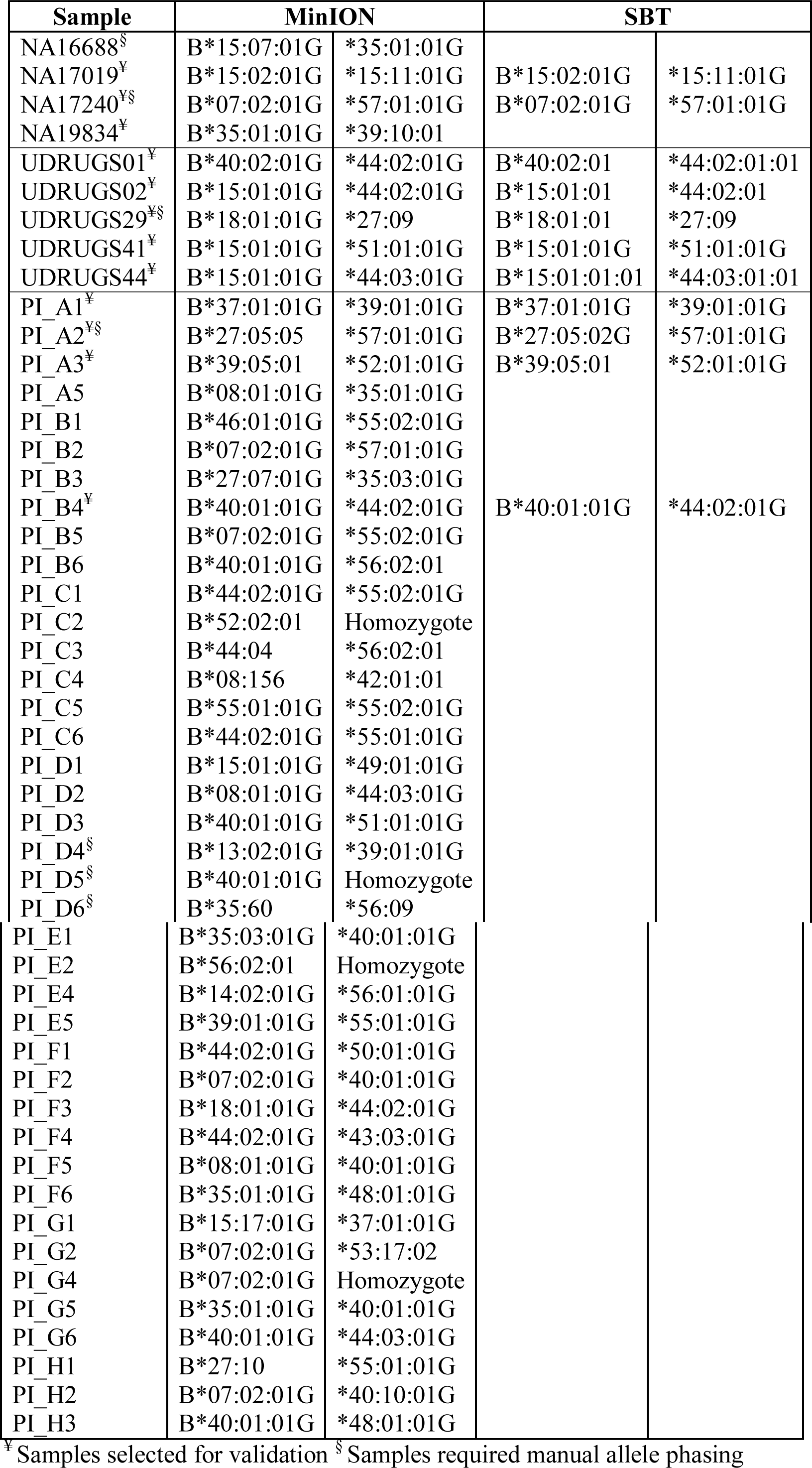
Assignment result of HLA-B obtained from MinION sequencing and SBT

| Sample | MinION |  | SBT |  |
| --- | --- | --- | --- | --- |
| NA16688<sup>§</sup> | B*15:07:01G | *35:01:01G |  |  |
| NA17019<sup>¥</sup> | B*15:02:01G | *15:11:01G | B*15:02:01G | *15:11:01G |
| NA17240<sup>¥§</sup> | B*07:02:01G | *57:01:01G | B*07:02:01G | *57:01:01G |
| NA19834<sup>¥</sup> | B*35:01:01G | *39:10:01 |  |  |
| UDRUGS01<sup>¥</sup> | B*40:02:01G | *44:02:01G | B*40:02:01 | *44:02:01:01 |
| UDRUGS02<sup>¥</sup> | B*15:01:01G | *44:02:01G | B*15:01:01 | *44:02:01 |
| UDRUGS29<sup>¥§</sup> | B*18:01:01G | *27:09 | B*18:01:01 | *27:09 |
| UDRUGS41<sup>¥</sup> | B*15:01:01G | *51:01:01G | B*15:01:01G | *51:01:01G |
| UDRUGS44<sup>¥</sup> | B*15:01:01G | *44:03:01G | B*15:01:01:01 | *44:03:01:01 |
| PI_A1<sup>¥</sup> | B*37:01:01G | *39:01:01G | B*37:01:01G | *39:01:01G |
| PI_A2<sup>¥§</sup> | B*27:05:05 | *57:01:01G | B*27:05:02G | *57:01:01G |
| PI_A3<sup>¥</sup> | B*39:05:01 | *52:01:01G | B*39:05:01 | *52:01:01G |
| PI_A5 | B*08:01:01G | *35:01:01G |  |  |
| PI_B1 | B*46:01:01G | *55:02:01G |  |  |
| PI_B2 | B*07:02:01G | *57:01:01G |  |  |
| PI_B3 | B*27:07:01G | *35:03:01G |  |  |
| PI_B4<sup>¥</sup> | B*40:01:01G | *44:02:01G | B*40:01:01G | *44:02:01G |
| PI_B5 | B*07:02:01G | *55:02:01G |  |  |
| PI_B6 | B*40:01:01G | *56:02:01 |  |  |
| PI_C1 | B*44:02:01G | *55:02:01G |  |  |
| PI_C2 | B*52:02:01 | Homozygote |  |  |
| PI_C3 | B*44:04 | *56:02:01 |  |  |
| PI_C4 | B*08:156 | *42:01:01 |  |  |
| PI_C5 | B*55:01:01G | *55:02:01G |  |  |
| PI_C6 | B*44:02:01G | *55:01:01G |  |  |
| PI_D1 | B*15:01:01G | *49:01:01G |  |  |
| PI_D2 | B*08:01:01G | *44:03:01G |  |  |
| PI_D3 | B*40:01:01G | *51:01:01G |  |  |
| PI_D4<sup>§</sup> | B*13:02:01G | *39:01:01G |  |  |
| PI_D5<sup>§</sup> | B*40:01:01G | Homozygote |  |  |
| PI_D6<sup>§</sup> | B*35:60 | *56:09 |  |  |

Table 3 (continued)
|  |  |  |
| --- | --- | --- |
| PI_E1 | B*35:03:01G | *40:01:01G |
| PI_E2 | B*56:02:01 | Homozygote |
| PI_E4 | B*14:02:01G | *56:01:01G |
| PI_E5 | B*39:01:01G | *55:01:01G |
| PI_F1 | B*44:02:01G | *50:01:01G |
| PI_F2 | B*07:02:01G | *40:01:01G |
| PI_F3 | B*18:01:01G | *44:02:01G |
| PI_F4 | B*44:02:01G | *43:03:01G |
| PI_F5 | B*08:01:01G | *40:01:01G |
| PI_F6 | B*35:01:01G | *48:01:01G |
| PI_G1 | B*15:17:01G | *37:01:01G |
| PI_G2 | B*07:02:01G | *53:17:02 |
| PI_G4 | B*07:02:01G | Homozygote |
| PI_G5 | B*35:01:01G | *40:01:01G |
| PI_G6 | B*40:01:01G | *44:03:01G |
| PI_H1 | B*27:10 | *55:01:01G |
| PI_H2 | B*07:02:01G | *40:10:01G |
| PI_H3 | B*40:01:01G | *48:01:01G |

\* Samples selected for validation § Samples required manual allele phasing

To validate this approach, we selected 11 individuals for HLA-B genotyping using the Sanger SBT method. These were four Polynesian, five UDRUGS and two Coriell samples. Six amplicons covering exon 2 and 3 of each individual were directly sequenced. HLA-B alleles were defined based on these polymorphic sequences. Using SBTengine (GenDX, Utrecht, Holland) for analysis, we found that there was a high consistency of variant calls compared with calls derived from the MinION data. Our genotyping data for all four Coriell individuals were concordant with the 1000 Genomes project. These results indicate that this MinION sequencing method is able to generate consensus sequences for high-resolution HLA-B typing with considerable accuracy.

## Discussion

There were 38 HLA-B alleles identified in the 40 Māori and Pacific Island individuals examined. Among these alleles, HLA-B∗40:01:01 had the highest frequency (28.95%), followed by HLA-B ∗44:02:01 (21.05%) and HLA-B∗07:02:01 (18.42%). According to the HLA allele frequency database (http://www.allelefrequencies.net), HLA-B∗40:01:01 is the most prevalent allele in the Han Chinese population (allele frequency of 0.155). It has been reported that the Polynesian people are ancestrally related to Micronesia, Taiwanese Aborigines and East Asia (Edinur *et al.*, 2013, Kayser *et al.*, 2008). This is consistent with our observation on the frequency of HLA-B∗40:01:01 in this population. On the other hand, the HLA-B∗44:02:01 and HLA-B∗07:02:01 alleles are present in multiple populations. When comparing with other studies on Polynesians, we found there were four alleles missing in our data (Edinur *et al.*, 2013).

Presumably this is because our sample size is insufficient to reflect the full range of HLA-B alleles in Polynesian or Māori populations. Nevertheless, the most common allele (HLA-B∗40:01:01) in our study is similar to the Edinur et al, findings, suggesting that this is a common HLA-B allele in this population. Interestingly, there was no HLA-B∗ 15:02:01 nor B∗58:01 observed in the Polynesian individuals. The lack of HLA-B∗58:01 in Polynesians has previously been reported in other studies (Abbott *et al.*, 2006, Roberts *et al.*, 2015). These alleles have been implicated in carbamazepine and allopurinol-induced severe cutaneous adverse reactions, respectively. However, HLA-B∗57:01:01, important for abacavir-associated hypersensitivity reactions, was apparent in two samples.

The primary goal of our study was to develop methods for HLA-B class 1 typing on the MinION nanopore sequencer. This study applied PCR across HLA-B exon 2 and exon 3, followed by barcoding and nanopore sequencing of 49 samples simultaneously. The high quality and good depth of coverage of our sequencing data for all samples, including several that were present at low concentration, enabled accurate assignment of HLA-B alleles. With ongoing improvements to the speed, throughput and workflow of MinION flowcells and the associated chemistry, it is likely that multiplexing could be extended to much greater numbers without compromising the ability for accurate typing.

The workflow we employed was straightforward and solely PCR-based. Though the ONT protocol requires 2.0 nM of input amplicon prior to the barcoding step, sample PI_G2 was successfully amplified and produced sequencing data at 0.19 nM, less than ten-fold the recommended amount. That said, even these poorly represented samples generated sufficient data for HLA-B typing. Varied number of reads between samples were observed, ranging from 80 to 17,329 reads. Despite imbalance issue, adequate depth of coverage was achieved for all samples and allowed for accurate HLA-B phasing. It is also worth noting that individual PI_C3 only had 80 reads that were aligned with reference sequences, but alleles could be assigned confidently. The ability to confidently call the genotype with such a small number of reads suggests that it would be reasonable to increase throughput by sequencing additional HLA loci, or indexing up to 96 individuals per sequencing run.

Our average read length was 1028 bp showing that our reads were long enough to cover exon 2 and exon 3 of HLA-B. We are aware that ambiguous typing of HLA-B may occur due to variants outside the region analyzed. As not all alleles have been completely sequenced, only exon sequences can be mapped in some cases. For example, there are 4,356 HLA-B alleles but only 384 alleles have complete sequence information (Robinson *et al.*, 2014). By this, we mean it is more practical to focus on exons solely than to sequence the entire gene for HLA-B typing with minimal ambiguity. Obviously, we are able to increase the read length for complete sequencing of the HLA-B if necessary, as the MinION is capable of very long sequence reads (Carter & Hussain, 2017).

Our results showed that SeqPilot was able to identify HLA-B alleles accurately using the MinION reads. Although the software could not automatically assign HLA-B alleles of all participants, it listed the most likely genotype combinations in order. This enabled us to re-phase the ambiguous positions and manually assign HLA-B alleles. Notably, the phasing bias mostly happened at nucleotide position 133-135 of exon 2. We realized that all these alleles were A/G heterozygous at nucleotide 133 and the sequence around this position was a repeat of G and A (GAGAG<u>R</u>GGAG). We also observed that if nucleotide 133 was G, there was usually a deletion of two to four nucleotides (Figure 3). It may be that deletion errors produced by the MinION combined with the complex nature of the HLA-B, make accurate interpretation of genotype particularly difficult around this region. Of the six individuals that were manually phased, alleles from three had been identified either by the SBT method or the 1000 Genomes project or both. These alleles were all consistent with our assignments.

The second aim of our study was to examine HLA-B alleles in individuals of Māori and Pacific Island descent living in New Zealand. Previous studies used allele-specific primer PCR for HLA-B genotyping, which provides low-resolution typing (Roberts *et al.*, 2013, Edinur *et al.*, 2013). Here, we report a feasible method for high-throughput and high-resolution HLA-B typing using NGS. Though ours is a relatively small sample, this initial finding can be used as a reference for future studies on the prevalence of HLA-B in these ethnic groups.

In conclusion, we have developed and evaluated a PCR-based HLA-B sequencing method using MinION Nanopore Technology. We have demonstrated our method is relatively straightforward and can generate accurate sequencing data from many barcoded samples in a single run. We also reported that precise HLA-B alleles could be obtained from the MinION reads with minimal phase ambiguity. Our protocol can be easily adapted for other HLA loci, or for full gene sequencing, or to employ greater levels of multiplexing. The method could be valuable in either a clinical or research setting.

## Acknowledgements

Kim Ton was supported by a University of Otago PhD scholarship. This work was supported by funding from the Jim and Mary Carney Charitable Trust.

